# A global survey of host, aquatic, and soil microbiomes reveals shared abundance and genomic features between bacterial and fungal generalists

**DOI:** 10.1101/2022.11.15.515575

**Authors:** Daniel Loos, Ailton Pereira da Costa Filho, Bas E. Dutilh, Amelia E. Barber, Gianni Panagiotou

## Abstract

Environmental change coupled with alteration in human lifestyles are profoundly impacting the microbial communities that play critical roles in the health of the Earth and its inhabitants. To identify bacteria and fungi that are resistant and susceptible to habitat changes respectively, we retrieved paired 16S and ITS rRNA amplicon sequence data from 1,580 host, soil, and aquatic samples and explored the ecological patterns of the thousands of detected bacterial and fungal genera. Through this large-scale analysis, we identified 48 bacterial and 4 fungal genera that were prevalent and abundant across the three biomes, demonstrating their fitness in diverse environmental conditions. The presence of generalists significantly contributed to the alpha diversity of their respective kingdom. Their distribution across samples explained a large percentage of the variation in the cross-kingdom community structure. We also found that the genomes of these generalists were larger and encoded more secondary metabolism and antimicrobial resistance genes, illuminating how they can dominate diverse microbial communities. Conversely, 30 bacterial and 19 fungal genera were only found in a single habitat, suggesting they cannot readily adapt to different and changing environments. These findings can contribute to our understanding of microbial niche breadth and its consequences for global diversity loss.

## Introduction

Environments, plants, and animals are colonized with communities of microbial organisms, termed the microbiome, which play critical roles in the function and health of their hosts and habitats. While understudied relative to bacteria, fungi play critical roles in environmental and host microbial communities, including important roles in carbon cycling and beneficial symbioses with plant and hosts (1–3). Importantly, environmental change and alterations in host lifestyle are profoundly affecting microbial consortia. Westernized diets low in fiber and rich in saturated fats and sugars, have decreased the abundance of beneficial microbes and been linked with myriad health conditions, including obesity, type 2 diabetes, and inflammatory bowel disease (4–7). Changes in marine environments due to climate change have induced major shifts in marine food webs, primary productivity, and carbon export (8–11). Additionally, anthropogenic climate change is resulting in net carbon loss in soil and changes in microbial community composition (12).

Ecological theory predicts that generalists, or organisms that are fit across a wider range of conditions, will be more resilient to changing environmental conditions (13, 14). Conversely, specialists, or organisms that are adapted to thrive in very specific environments, will be less able to withstand perturbations to their habitat. As the Earth and its inhabitants are experiencing unprecedented changes to their health and habitats, it is crucial to understand the capacity of individual microbial taxa to adapt to changing environmental conditions. Those unable to change are susceptible to biodiversity loss, while generalists that can grow in a wider range of conditions may survive and flourish with unknown consequences. Meta-analyses of bacterial community data have identified ecological and evolutionary features of bacterial generalists and specialists (15, 16). However, a corresponding study examining these features in fungi has not been performed. Moreover, bacterial and fungal kingdoms have not been considered together at the global scale, despite substantial evidence from individual settings that bacteria and fungi commonly interact with each other with pronounced consequences (17, 18).

To this end, we performed a large-scale analysis of community sequencing data sets from host, soil, and aquatic environments with paired bacterial and fungal characterization to shed light on the ecological properties of the genera present and their putative resilience to change. We focused on three aspects: (i) the identification of bacteria and fungi that occurred in diverse environments capable of adapting to diverse environments (generalists) or were limited to highly specific environments (specialists); (ii) the relative abundance of bacterial and fungal generalists and specialists as a marker for their fitness and competitive colonization potential; and (iii) whether their presence in a habitat was associated with global changes in inter- and cross-kingdom population structure.

## Results

### Environmental specificity of bacterial and fungal communities

For a global survey of bacteria and fungi across microbial communities, we analyzed paired 16S and ITS rRNA amplicon sequence data from 1,580 samples deposited in public databases. Samples were collected from Europe, Asia, and the Americas between 2010 and 2018 (Figure 1A). For cross-biome comparisons, samples were classified as aquatic, host, or soil environments based on the habitat they were collected from. This broad grouping is supported by principle coordinate analysis (PCoA) based on Bray-Curtis dissimilarity showing that samples from each environment largely cluster with each other and distinct from the other environments (Figure 1B)—a finding mirrored by a recent study of 22,700 bacterial microbiomes (15).

**Figure 1:**
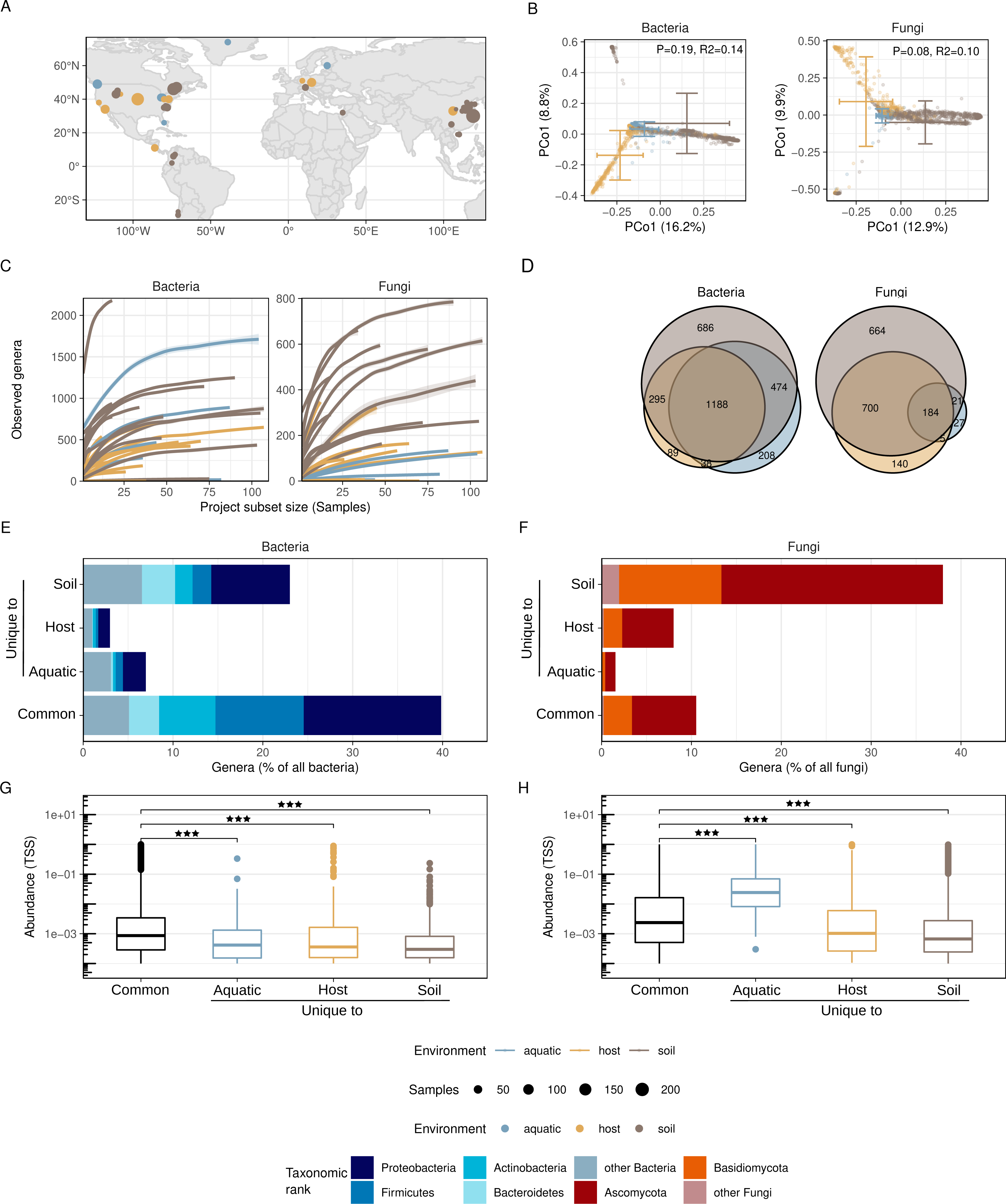
A global analysis of microbial communities reveals differences in environmental specificities between bacteria and fungi. (A) Distribution of samples used in this study (N=1,580) by geographic location. (B) Bray-Curtis dissimilarity between samples, colored by environment. Crosshatches represent the mean ± SD for each environment. Significance calculated by PERMANOVA. (C) Rarefaction curves of Shannon alpha diversity for each study demonstrate sufficient sampling depth. Curves are shown as LOESS regressions from 10 independent sampling trials at 10 given sampling subset sizes. Lines are colored by environment and are surrounded by ribbons indicating the 95% confidence interval across the trails. (D) Intersection of bacterial and fungal genera found in at least one sample in each environment as Venn diagrams. (E,F) Percentage of genera found in all three or only one environment. (G,H) Abundance comparisons of common and unique genera by total sum scaling (TSS). A genus was considered present in a sample using a threshold of abundance > .01%. Significance determined by Wilcoxon rank sum test; *** denotes p < 0.001.

Of the 1,580 samples that we analyzed, 871 originated from soils, 494 from hosts (including mammalian, non-mammalian, and plant hosts), and 215 from aquatic environments. The habitats that contributed the largest number of samples for each environment were temperate (N=498) and conifer forests (N=147) for the soil, gut (N=287) and skin (N=68) for the hosts, and large lakes (N=87) and other freshwater (N=71) for the aquatic environments (see Supp. File 1 for details of all projects). Due to the limited availability of fungal community data in marine samples, this large habitat was only represented with 10 samples. Taxonomic profiling of bacterial and fungal communities was performed using the SILVA and UNITE databases, respectively. Rarefaction curves of each habitat indicated that most projects adequately captured the diversity of both the bacterial and fungal communities (Figure 1C). In total 2,977 bacterial and 1,740 fungal genera were detected across all samples (Figure 1D). We next examined the overlap of genera between environments, where a genus was considered shared if it was detected in at least one habitat in each of the three different environments (host, aquatic, and soil). For bacteria, soil and aquatic environments had the highest number of shared genera (N=1,662), followed by host-soil (N=1,483) and host-aquatic (1,226). The pattern was different for fungi, with host-soil sharing the most (N=884), followed by aquatic-soil (N=205) and host-aquatic (N=189). These trends remained after controlling for the different number of samples across the three environments in 842 and 998 out of 1,000 random down sampled subsets for bacteria and fungi, respectively. Finally, we also confirmed that a similar degree of overlap between the environments was observed for different 16S and ITS amplicons, as well as significant correlations in the abundances of individual genera (Supp. Figure 1A-C).

While 40% of the total bacterial genera were found in all three environments, the percentage dropped to only 11% for fungal genera, indicating a higher degree of environmental specificity (Figure 1E & F). The most prevalent higher order taxonomic ranks that were detected in all three environments were *Proteobacteria* followed closely by *Firmicutes* for bacteria, and *Ascomycota* for fungi. For both bacteria and fungi, soil was the environment with the highest percentage of uniquely detected genera (i.e. genera not detected in any sample from host or aquatic origin), with 23% and 38%, respectively for each kingdom. While aquatic-specific bacteria accounted for 7% of the total number of detected genera, the percentage of unique fungi in aquatic samples was only 2% (Figure 1 E & F). The opposite trend was observed for host-associated microbes, with only 3% and 8% of unique bacteria and fungi, respectively in this environment.

We subsequently compared the relative abundance of genera that were found in all environ-ments or were uniquely detected in soil-, host-or aquatic-associated environments. Bacterial genera detected in all three environments were significantly more abundant (Wilcoxon Rank Sum test, p < 0.001) than genera uniquely detected in one of the environments (Figure 1G). A similar pattern was observed with fungi. However, a notable exception was the relatively high abundance of fungi that were uniquely detected in aquatic samples. Genera of aquatic fungi were more abundant than either common genera and uniquely detected in soil-or host-associated environments (Figure 1H). This observation was also robust across the different 16S and ITS regions used in the dataset (Supp. Figure 1D). Taken together, we find that soil bacteria and fungi show a higher degree of biome specificity compared to aquatic and host environments, and that genera detected in all three environments were also more abundant than those only detected in only one environment.

### Bacterial and fungal generalists are more abundant than specialists and have distinct genomic features

Generalists and specialists play important, yet distinct roles in ecosystems. However, objectively identifying them has proven challenging. To define multi-kingdom generalists and specialists, we set the following criteria: generalists are genera found with high prevalence (>40%) in at least one habitat (e.g. gut, boreal forest) from each of the three environments (host, aquatic, soil), where prevalence was defined as a relative abundance above 0.01% in >1 sample from the habitat. Conversely, specialists were genera with a high prevalence (>40%) in one habitat and low prevalence (<5%) in every other. Using this approach, we detected 48 bacterial generalists and 30 specialists (Supp. Table 1). To ensure that the generalists identified were legitimately present in the microbial community and not the result of reagent or sequencing contamination, we performed decontamination on low biomass projects, including all aquatic projects and low biomass host sites like the lung (see Methods for details). There were no significant differences in the abundance the generalists before and after sequence data decontamination (Supp. Figure 2). As an additional control, we compared the DNA extraction methods/kits used, any DNA purification kits utilized, library preparation kit, and the sequencing facility of the projects included in our study. We could not find appreciable overlap among the project studies that would imply that the generalists identified were the result of contamination via common methodology or reagents (Supp. File 2).

To confirm our definition of generalists and specialists, we calculated Levins’ niche breadth indices (B_n_), which measures taxon distribution across environments and where higher values indicate even distribution across environments (19). Generalists showed significantly higher B_n_ values than specialists (Wilcoxon Rank Sum test, p < 0.001; Supp. Figure 3A). All specialists and all generalists, with the exception of the *Christensenellaceae R7* group, were above the detection limit and had a significant Levins’ niche breadth signal after Benjamini-Hochberg adjustment (19). As our criteria for defining generalists and specialists was reliant on human-defined biome annotations, we further validated our approach by comparing it to the recently developed social niche breadth (SNB) score (15). By comparing the similarity or diversity of microbial communities where a given genus occurs, SNB provides a data-driven score independent of biome annotations based on an independent dataset of over 22,500 bacterial microbiomes (15). Indeed, the generalists identified in our study had significantly higher SNB scores than the bacterial specialists we identified (Wilcoxon Rank Sum test, p < 0.001; Supp. Figure 3B).

We observed multiple phylogenetic origins for both generalists or specialists (Chi squared test, p > 0.05), indicating their roles as generalists and specialists evolved independently (Supp. Figure 4). Each of the top five bacterial generalists were detected in more than 50% of the 1,580 samples. Among them, the most prevalent was *Pseudomonas*, which was detected in 52%, 70% and 89% of host, soil and aquatic samples respectively, followed by *Bacillus* (33%, 71%, 35%) and *Bradyrhizobium* (17%, 73%, 35%). The most extreme bacterial specialists came from the soil. While *Gryllotalpicola* and *Anaerovibrio* were found in >91% of biochar samples, the prevalence dropped to 0.1% on average for non-soil environments (Supp. Table 1). Specialists were also found in host- and aquatic-associated environments. For example, *Acetatifactor* was found in 80% of samples from the murine gut, but had a prevalence of <3% in all other habitats. The genus *Leptospira* was found in 83% of samples of the Cuyahoga River, but had a prevalence less than 2% in all other habitats. Interestingly, when comparing the relative abundance of generalists and specialists, we observed that both bacterial and fungal generalists had a significantly higher abundance (Figure 2A, Wilcoxon Rank Sum test, p < 0.001). This pattern remained when we used stricter and looser thresholds to define generalists and specialists (Supp. Figure 5). This finding confirms the pattern observed above (Figure 1G, H), suggesting that independently of how groups are defined, genera that can colonize diverse environments are usually able to outcompete habitat-specific genera.

**Figure 2:**
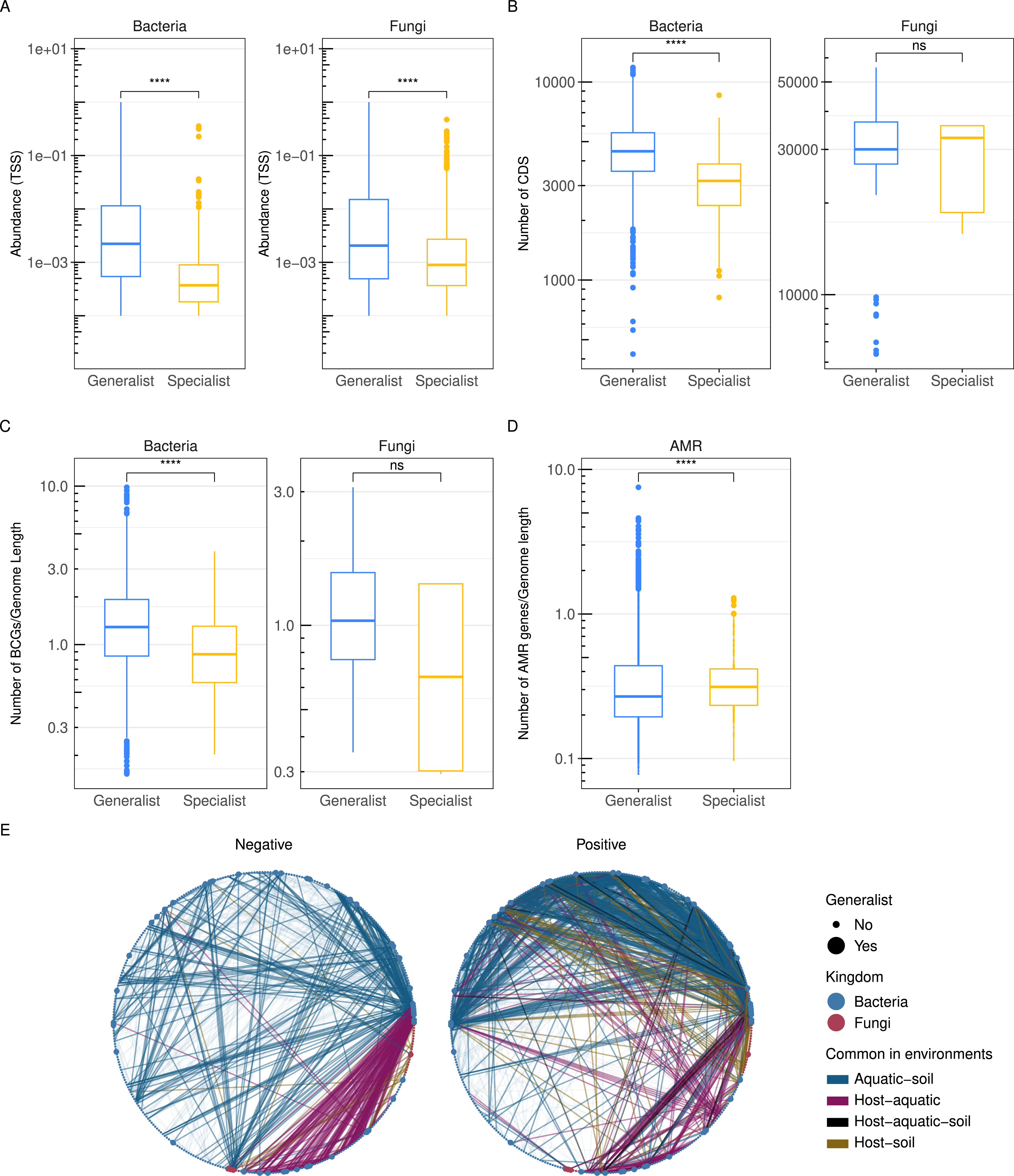
Generalists are more abundant and bacterial generalists have larger genomes with more biosynthetic gene clusters and antimicrobial resistance genes. (A) Relative abundances of bacterial and fungal generalists and specialists. Values were averaged by project to account for different cohort sizes. Statistical significance calculated using Wilcox rank-sum test (**** denotes p<0.0001). (B-C) Number of coding sequences (CDS) (B) and biosynthetic gene clusters (BCGs) (C), in the genomes of generalists and specialists normalized to their genome length in bp. Data from the genomes of 2,328 bacterial generalists, 117 fungal generalists, 471 bacterial specialists, and 5 fungal specialists. Statistical significance calculated by Wilcox rank-sum test (** denotes p<0.01, *** denotes p<0.001; **** denotes p<0.0001; ns denotes p>0.05). (D) Number of antimicrobial resistance (AMR) in the genomes of bacterial specialists, normalized by genome length. (E) Networks of genera found in all three environments and significantly co-abundant in the majority of environments (SparCC FDR p < 0.05, |r| > 0.2).

When looking at the fungal kingdom, the number of generalists was much lower and only *Aspergillus*, *Malassezia*, *Aureobasidium*, and *Cortinarius* satisfied the criteria of a generalist (Supp. Table 1). Among these, *Aspergillus* had the highest overall prevalence among all samples with 38%, 52% and 12% in the host, soil, and aquatic samples, respectively. From the 19 fungal specialists, *Chrysanthotrichum* and *Mycocentrospora* were the most habitat-specific, with prevalences of 68% and 48% in temperate and conifer forests, respectively, but a mean prevalence of 0.1% in all other habitats. Only two of the 19 fungal specialists (11%) originated from outside soil environments (*Vuilleminia* and *Seimatosporium* from plants). As with bacterial genera, the relative abundance of fungal generalists was significantly higher than that of fungal specialists (Figure 2A, Wilcoxon Rank Sum test, p < 0.001).

To gain insight into how generalists achieve high abundance in diverse environments, we analyzed the genomes of generalists and specialist genera available on NCBI (see Methods for details on genome selection). For bacteria, when analyzing the genomes of 2,328 generalists and 471 specialists (Supp. File 3), the generalists had significantly larger genomes, as measured by the number of coding sequences (CDS) with a mean of 4,671 CDS for generalists and 3,189 for specialists. (Wilcoxon Rank Sum test, p < 0.001; Supp. File 3). This trend remained after controlling for genome length with generalists having a mean of 925 CDS/Mb genome compared to 921 CDS/Mb for specialists (Wilcoxon Rank Sum test, p = 0.013). As secondary metabolism genes are often used by microbes during competition for resources and as chemical warfare in crowded environments, we examined the genomes of generalists and specialists for the presence of biosynthetic gene clusters (BCGs). Strikingly, the genomes of bacterial generalists encoded significantly more BCGs with an average of 7.4 BGCs and 1.4 BGCs/Mb genome compared to 2.7 BGCs and 1.2 BGCs/Mb for specialists (Figure 2C, Wilcoxon Rank Sum test, p < 0.001). Further differentiating bacterial generalists, they also contained significantly more antimicrobial (AMR) genes, with an average of 4.2 AMR genes and 0.39 AMR genes/Mb genome compared to 1.0 AMR genes and 0.36 AMR genes/Mb for specialists (Figure 2D; Wilcoxon Rank Sum test, p < 0.001). For fungi, no significant differences in either the number of CDS or BGCs was observed (Figure 2B, C), likely due to the severe underrepresentation of publicly available fungal specialist genomes (N=5).

To explore intra and inter-kingdom interaction patterns and to gain further insight into the downstream effects of the observed differences between generalists and specialists, we constructed individual co-abundance networks for soil, host, and aquatic environments (see Methods for details). Despite only considering the 1,188 bacterial and 184 fungal genera commonly detected in all three environments, the topological characteristics of the networks for each environment were highly distinct, as measured by significant differences in betweenness and Kleinberg’s hub node centrality scores (Wilcoxon test, p < 0.001; Supp. Figure 6). In spite of the differences in topology, we could still compile subnetworks of inter- and intra-kingdom correlations found jointly among host-soil, host-aquatic and/or soil-aquatic environments. Strikingly, 45 of the 48 bacterial generalists and all 4 fungal generalists were part of those subnetworks, which are characterized by a higher number of positive than negative edges (Figure 2E). The ratio of positive to negative edges was higher in correlations involving a generalist (2.5) compared to all other edges (2.2). When we looked for interactions between genera found in all three environments, we identified 43 such edges that all represented positive interactions between bacteria and included 21 generalists. Together, these findings suggest that the success of generalists in colonizing diverse environments and achieving high abundances may be attributable to their ability to carve out a niche for themselves using secondary metabolism and AMR genes, and by eliciting positive interactions with other highly prevalent genera.

### Bacterial generalists exert a strong influence on the intra- and inter-kingdom community structure

We subsequently explored whether the presence of generalists and specialists had an impact on the diversity of a community. Interestingly, alpha diversity, as measured as Chao 1 and Shannon, was significantly lower in samples where no generalist was detected compared to samples with generalists present for both bacterial (Figure 3A) and fungal (Figure 3B) communities (Permutation test of samples lacking any of the N generalists compared with samples lacking any N random taxa, 1*10^4^ permutations, p < 0.03). Conversely, the impact of specialists on alpha diversity in their specific habitat was much less profound and varied by habitat without a clear trend (Supp. Figure 7).

**Figure 3:**
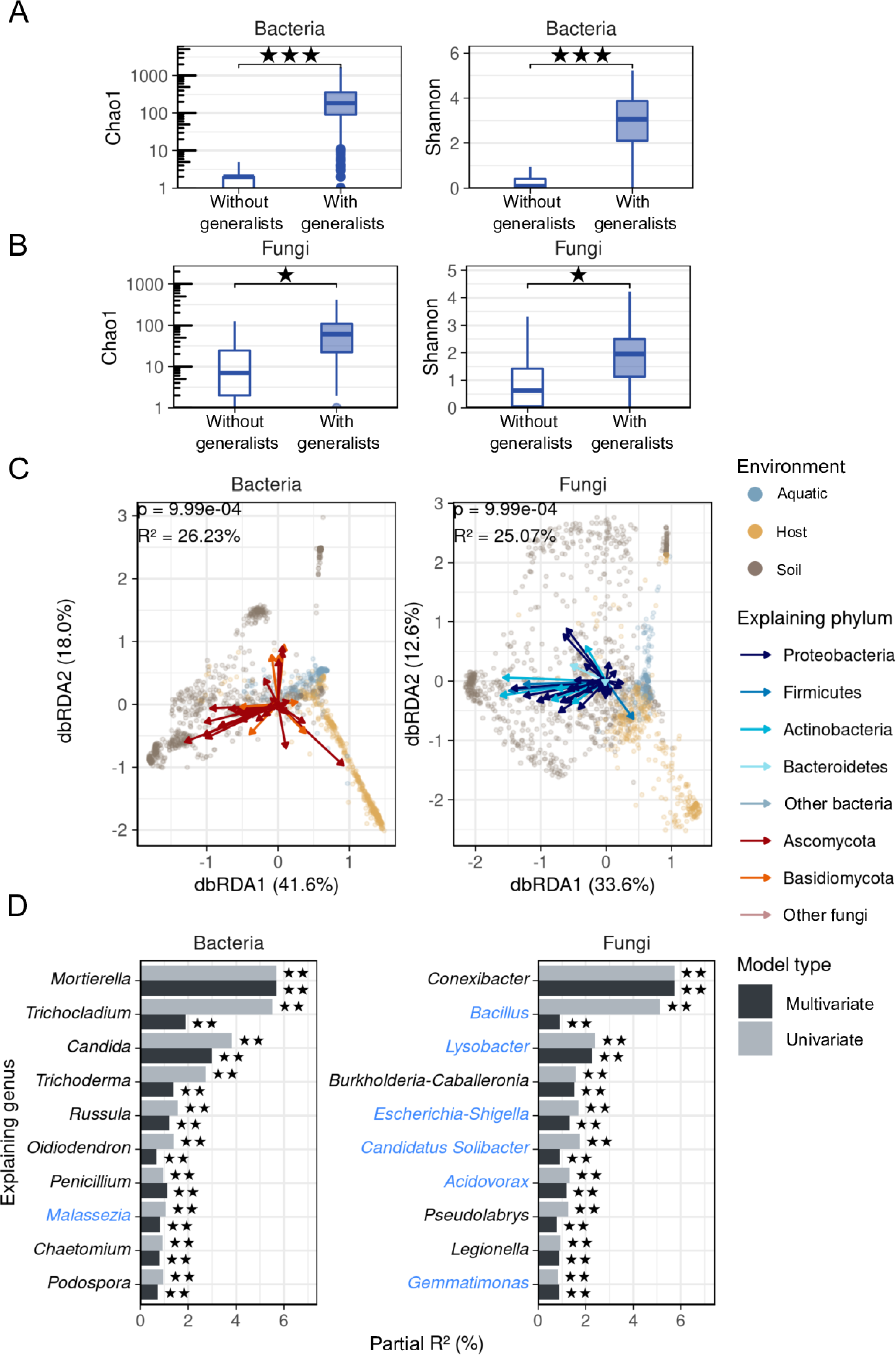
Generalists significantly impact diversity and cross-kingdom variation. A-B: Shannon and Chao1 alpha diversity were calculated for bacteria (A) and fungi (B). Samples were grouped by whether they contained any generalist (genera with >40% prevalence in at least one habitat from every environment; abundance > 0.01%; N=1,500 for bacteria, N=1125 for fungi), or do not contain any generalist (N=91 for bacteria; N=466 for fungi). Significance bars indicate permutation test compared to samples without random taxa instead of generalists (* q<0.05, ** q<0.01, *** q<0.001). C-D: Bacterial and fungal Bray-Curtis dissimilarities constrained by explanatory genus abundances of the other kingdom using distance-based Redundancy Analysis (dbRDA). C: Explaining genera were selected using a feedforward approach. Effect size of most explanatory taxa is shown in D by multivariate model (as displayed in C) or univariate model containing only the taxon of interest. Generalists are indicated with blue text. Stars indicate significance by ANOVA (** p<0.01).

We subsequently shifted our focus to inter-kingdom interactions which are often overlooked in microbial ecology studies and examined bacterial generalists for a role in shaping the myco-biome community structure and vice versa. As expected, we observed a significant separation between the soil, host, and aquatic micro- and mycobiome beta diversity by Bray-Curtis dissimilarity (Figure 3C, PERMANOVA, p < 0.001 for both bacteria and fungi). Constrained ordination revealed a significant, linear relationship between bacterial Bray-Curtis dissimilarity and fungal community composition and vice versa (Distance-based redundancy analysis, abbreviated dbRDA, ANOVA p < 0.04 for all explanatory genera of the other kingdom in a multivariate model). Bacterial genera could explain an extensive part of the mycobiome variation observed in the three environments with a partial R2 of 25% by dbRDA. Of the bacterial genera, *Conexibacter*, *Bacillus*, and *Lysobacter* had the highest explanatory power on mycobiome variation (Fig 3D). Interestingly, six out of the top ten explanatory bacterial genera in the dbRDA were generalists. Similarly, fungal genera explained 26% of the microbiome variation between host, soil, and aquatic samples, with *Mortierella*, *Trichocladium*, and *Candida* having the highest explanatory power (Fig 3D). Among the top ten explanatory genera was one of the four fungal generalists -*Malassezia*. Altogether, our analysis indicates that bacterial and fungal generalists profoundly impact microbial communities by contributing positively to the taxonomic (alpha) diversity of their kingdom-an ecological characteristic often associated with healthy environments, and they can also contribute to shaping cross-kingdom microbial structures.

## Discussion

Recent global changes are profoundly affecting the health of our planet and its inhabitants (20–22). As environmental and host-associated microbial communities are increasingly exposed to changing habitats, we still lack knowledge about the capacity of millions of bacterial and fungal species to cope with these shifts. With this in mind, we performed a large-scale global survey of host, aquatic, and soil microbiomes to reveal ecological and genomic properties of bacterial and fungal genera that may promote or limit their establishment in new environments and how they contribute to the richness and diversity of an environment. Analysis of 1,580 paired host, soil, and aquatic micro- and mycobiomes identified ∼70 specialist genera whose limited distribution suggests they may struggle in different or changing habitats and identify ∼50 widely distributed genera with a clear ability to thrive in many environments. Through this analysis, we also identified genomic and ecological properties associated with generalists, including their contributions to alpha diversity, and structuring the beta diversity in the other kingdom. Generalists also had larger genomes with more secondary metabolism genes, suggesting a mechanism for how they thrive in such diverse and often highly competitive habitats.

While the concept of generalists and specialists is not new to ecology, it has mostly been applied in specific habitats (23–27) and not on a global scale. Similarly, while studies of generalist and specialist microbes have been carried out (15, 16, 24, 28–32), they have rarely considered eukaryotic microorganisms such as fungi, despite the critical role fungi play in many habits (33–38). As a result, the biotic interactions that shape microbial communities across kingdoms has not been investigated on a global scale. We demonstrate that both bacterial and fungal generalists share ecological features, including the ability to reach significantly higher abundances than specialists and contributing positively to the richness and diversity of their respective kingdoms. Moreover, six bacterial generalists, including *Bacillus*, *Lysobacter*, *Escherichia* and *Gemmatimonas* and one fungal generalist, *Malassezia*, harbor additional ecological properties and appear to play a significant role in shaping cross-kingdom microbial composition (Figure 3D).

These positive roles for generalists are somewhat at odds with previous work showing that generalists negatively impact ecosystems through homogenization (39–41). One possible explanation is that the species- and strain-level diversity of the microbial world is enormous compared to higher eukaryotes, where many key studies have been conducted. The variable and pronounced effects of strain- and species-level diversity within generalist genera are highlighted in biocontrol agents. Strains of the generalist bacteria *Bacillus*, *Pseudomonas*, and *Streptomyces* are approved as Biocontrol agents for soil-borne diseases in the European Union (Source: EU Active Substance Pesticide Database, accessed December 2023). However, other strains of *Bacillus* and *Pseudomonas* are pathogens of crops.

Indeed, a challenge for the future will be to move the analysis of microbial generalists and specialists beyond taxonomic description to understanding the functional characteristics that distinguish them from other taxa. Our study was carried out at the genus level due to the limitations of accurate species-level classification of bacteria and fungi via amplicon metabarcoding (42–44). For bacteria, one way forward may be deep functional characterization at the pathway and enzyme level using shotgun metagenomics datasets. The functional characterization of fungal generalists and specialists will prove to be a much greater challenge. Their genomes are larger and more complex, and their physiology less studied. Consequently, functional prediction tools and community-level modeling based on metagenomic data for fungi lags behind those for prokaryotic microorganisms.

Overall, the generalists and specialists identified cumulatively account for a small fraction of the total taxonomic diversity of each kingdom (<2.6% of total bacterial genera and <1.3% of fungal genera). However, we find that generalism is proportionally rarer in fungi, with 1.6% of bacterial genera in the dataset meeting the definition for generalism compared to 0.2% of fungi. The reason for this is an intriguing question for further work, but we posit that it may be due to differences in the genome dynamics between bacteria and fungi. Bacterial genomes are more flexible and shaped by horizontal gene transfer to a degree not seen in fungi. Alternatively, differences in dispersal may allow for bacterial genera to expand to different habitats more efficiently than fungi.

Our finding that bacterial and fungal generalists have larger genomes is consistent with the expectation that fitness across diverse habitats requires a larger genomic repertoire and parallels the results from another meta-study of bacterial community data (16). In our work, we were able to delineate some of the specific tools used by generalists, such as the acquisition of an enrichment of biosynthetic gene clusters and antimicrobial resistance genes. However, it is likely that other genomic factors underlie the generalist ability; the relative contributions of these factors involved in inter-microbe communication, compared to other genomic features such as metabolic versatility, is an open question.

One limitation of our study is that aquatic, and particularly marine, fungi are understudied. This led to the exclusion of important marine habitats, such as seawater and marine sediment from our dataset, due to the lack of paired, publicly available 16S-ITS samples at the time we assembled our dataset. When we analyzed several marine projects *post hoc* representing seawater, marine sediment, and coastal seagrass water (45–47) we did not iden-tify any bacteria that were not already classified as generalists or specialists compared to our original dataset. On the other hand, we identified *Puccinia* as a novel fungal marine specialist and *Cladosporium*, *Penicillium*, and *Chateomonium* as previously unrecognized fungal generalists. The ecological and genomic features of these taxa will be interesting to examine in the future. An additional caveat is that the relationship between habitats, es-pecially between a host and its environment, is complex. For example, plants have selective effects on microbial diversity in the rhizosphere (48–50). Nevertheless, in our study and other global microbiome studies (15, 16), rhizosphere microbial communities more closely resemble soil than other host-associated habitats, such as the host-associated microbiomes of insects, birds, and mammals.

Finally, the observation that roughly a quarter of bacterial community composition could be explained by fungal abundance, and vice versa, strongly emphasizes the role of the multi-kingdom interactions in microbial communities and highlights the amount of information potentially missed by examining only one kingdom—an important point for future microbial community studies. In conclusion, our global survey of bacterial-fungal communities has generated a valuable list of genera that may be vulnerable to biodiversity decline and even extinction under changing environmental threats (51, 52). Conversely, the generalist bacteria and fungi identified are highly resilient against environmental perturbations. However, their functional roles in ecosystems, especially at the species- and strain level, will benefit from approaches that combine large-scale computational analyses and laboratory experiments. Together, these interdisciplinary approaches can address the many open questions about microbial niche range and its consequences for microbial extinction and global biodiversity loss.

## Author contributions

Conceptualization: A.E.B, G.P; Methodology/Validation: D.L, A.E.B, G.P, B.E.D; Visualization: D.L, A.P.d.C.F; Project Administration/Supervision: A.E.B, G.P; Funding Acquisition: A.E.B, G.P; Writing – Original Draft: D.L, A.P.d.C.F, A.E.B, G.P.; Writing – Review & Editing: all authors.

## ACKNOWLEDGMENTS

This work was funded by the Deutsche Forschungsgemeinschaft (DFG, German Research Foundation) under Germany’s Excellence Strategy – EXC 20151 – Project-ID 390813860. BED is supported by the European Research Council (ERC) Consolidator grant 865694 and the Alexander von Humboldt Foundation in the context of an Alexander von Humboldt-Professorship.

## Declaration of Interests

The authors declare no conflict of interests.

## Star Methods

### RESOURCE AVAILABILITY

#### Lead contact

Further information and requests for resources should be directed to and will be fulfilled by the lead contact, Amelia E. Barber or Gianni Panagiotou.

#### Materials availability

This study did not generate new unique reagents.

#### Data and code availability

This paper analyzes existing, publicly available data. These accession numbers for the datasets are listed in the Supplementary file 1. All original code has been deposited at https://github.com/bioinformatics-leibniz-hki/its-16s and is publicly available as of the date of publication.

## METHOD DETAILS

### Sample selection

Included studies were retrieved by querying NCBI BioProject with the terms ‘bacteria’ and ‘fungi’ in any field. Only BioSamples with both 16S rRNA and ITS amplicon sequencing data were considered for the concurrent analysis of both kingdoms. We also required that 16S and ITS sequences were deposited under a unified BioSample ID to definitely link patterns in bacterial and fungal diversity. This excluded some additional projects, as the 16S and ITS sequences were deposited under different BioSample IDs. We used both the identifier and attributes of the BioSample, such as aliases and library names, to map fungal and bacterial read files to a sample using a custom script. Samples were associated to an environment (aquatic, host, or soil) using manual curation of associated publications and BioSample attributes provided by the depositor. The three environments were further subdivided into 17 habitat groups based on the body part and/or the ecoregion of the sampling location for host and other samples, respectively (53). Habitats with less than five samples were pooled together.

### Generation of genus-level abundance profiles

Genus-level abundance profiles were calculated using a custom nextflow pipeline (54). Briefly, reads were downloaded from NCBI SRA using grabseqs, except for the American Gut Project, which was downloaded from Qiita (55, 56). Paired-end reads were merged using NGmerge (57). Quality Control (QC) and adapter removal was performed using trimmomatic with a minimum Phread quality of 20 and a minimal read length of 100 (58). Quality was assessed using FastQC and MultiQC (59). Subsequent steps were performed using QIIME2 (60). Reads were dereplicated following closed-reference OTU picking for both kingdoms separately using VSEARCH with a 97% identity threshold (61). For taxonomic annotation, SILVA 132 97% consensus and UNITE 8.2 dynamic databases were used for bacteria and fungi, respectively (62, 63).As detection of archaea was highly variable across the 16S datasets, any counts assigned to archaea were removed prior to downstream analyses. Relatedly, co-amplifying plant and non-fungal microbial eukaryote sequences were excluded from analysis as we used a version of the UNITE database that only included fungal sequences. Following quality control, a total of 1,580 samples were selected for downstream analyses.

### Discovery of sample rRNA amplified region

Multiple rRNA regions were used to characterize microbial diversity as the study dataset is composed of many sequencing projects. When available, the specific rRNA region amplified was obtained from deposited metadata or linked publication. For BioProjects where this information was not available, the following was performed. As the SILVA database (v138.1) contains full length bacterial rRNA sequence, the hypervariable regions (e.g. V1-V3, V4-V5) from each taxon was extracted using the in silico pcr tool (https://github.com/egonozer/in_silico_pcr) with primers described in (64).Amplicon sequence data from each project was then aligned to each variable region using BWA-MEM v.0.7 and contig coverage quantified using BBTools v.39.01. The 16S variable region with the highest percent coverage was taken as the region amplified in the study. For the ITS amplicon data, ITSx 1.1.13 (65) was used to extract the ITS1 and/or ITS2 consensus from sequence reads. The BioProject primers identified through this analysis, as well as those retrieved from association publications is listed in Supp. File 1.

### Abundance correlation between varying rRNA amplicons

To calculate the correlation in genus abundances between the differing rRNA regions amplified, genera that were detected in all three environments were considered and samples aggregated into whether they included sequence from the V1-V4 regions or V4-V5 regions for bacteria and ITS1 or ITS2 for fungi. For each rRNA category, Pearson’s correlation coefficients were calculated for genus abundance in each environment. The similarity between the correlation matrices (V1-V4 and V4-V5 for bacteria and ITS1 and ITS2 for fungi) was then calculated by transforming the upper triangle of each correlation matrix into a vector and calculating the correlation coefficient between the two.

### Decontamination of low biomass projects

Low biomass samples are susceptible to amplification of low-level contaminating sequences in extraction kits and other reagents. To control for this, low biomass projects, including all aquatic project and low biomass host habitats like the long were decontaminated using the decontam R package (66). The majority of low biomass projects contained blanks or negative controls and were decontaminated using the “prevalence” method of decontam. In this method, OTUs observed in the non-sample control samples are labeled as contaminants and removed from the abundance table. One low biomass project did not have sequencing controls but provided us with the input DNA concentrations used for library preparation, as determined by Qubit (Invitrogen). This allowed the samples from this project to be decontamined using the “frequency” mode of decontam. In this mode, contaminating OTUs are identified and removed from downstream analyses as their abundance inversely correlates with the input DNA concentration, rather than independent of it. The mode of deconam used for each project is indicated in Supp. Figure 2.

### Genome features of generalists and specialists

As amplicon sequence data is based on maker genes, deposited genomes were used to characterize functional traits associated with the genomes of generalists and specialists. The generalists and specialist genera were queried in NCBI RefSeq. Of the resulting genome list, all genomes or up to 60 randomly selected genomes if more were available were selected for each genus. This resulted in genomes for 2,328 bacterial generalists, 117 fungal generalists, 471 bacterial specialists, and 5 fungal specialists. Genome size and number of coding regions were obtained from the NCBI metadata. For the calculation of the number and type of biosynthetic gene clusters in each genome, AntiSMASH v6.1.1 was used (67). Antimicrobial and stress resistance genes were predicted in bacterial genomes using AMRFinderPlus (68). Counts were divided by genome length for normalization.

## QUANTIFICATION AND STATISTICAL ANALYSIS

### Workflow and statistical analysis

Analyses were performed using a custom drake pipeline (69) built using the programming language R 4.0.2. Briefly, abundances obtained from OTU profiling were total-sum-scaled (TSS) and pooled at genus rank. All tools were used with default parameters if not explicitly specified.

### Diversity

Alpha diversity was estimated using Shannon and Chao1 metrics with the phyloseq and vegan packages (70, 71). To quantify the contributions of a bacterial community profile with the fungal one and *vice versa*, we used linear and unsupervised Canonical Correlation Analysis, as implemented in the function CCorA of the vegan R package (70). P-values were obtained using blocked permutations to control for the habitat and to reduce assumptions of the test. Supervised constrained ordination was performed using stepwise Distance-based Redundancy Analysis (dbRDA) adapted from (72). This analysis shows linear relationships between bacterial dissimilarities and abundances of selected explanatory fungal genera (and *vice versa*). An optimal subset of up to 50 explanatory genera of the other kingdom was computed using a stepwise feed-forward approach, as implemented in the ordistep function of the vegan R package (70).

### Co-abundance networks

SparCC, as implemented in FastSpar, was used to assess correlation between taxa pairs for each environment separately (73, 74). Both kingdoms were pooled together, allowing for the identification of interkingdom correlations. Only genera found in all three environments were considered for pairwise correlation. Node topology metrics were calculated using the R package igraph.

### Generalists and specialists

Genera were defined as generalists if they were found in at least 40% of samples in at least one habitat from each environment (host, soil, aquatic) with a relative abundance of at least 0.01%. Conversely, genera were defined as specialists if they were found in at least 40% of samples in one habitat and less than 5% of samples in all other habitats using the same abundance threshold as for generalists. Levins’ niche breadth index was calculated as implemented in the R package MicroNiche (75). Social niche breadth (SNB) was calculated as in von Meijenfeldt et al., Nature Ecology and Evolution (2023) using the data from the MGNIFY database analyzed in this study.

## Supplementary file titles

Supplementary file 1: BioProject IDs and metadata for the datasets analyzed in this study.

Supplementary file 2: Kits used for DNA extraction, DNA purification, DNA amplification and library preparation by the sequencing projects included in this study.

Supplementary file 3: Accession numbers and genome characteristics for the gen-eralists and specialists profiled.

## Notes

### Competing Interest Statement

The authors have declared no competing interest.

### Summary of Updates

Added work ruling out reagent and sequence contamination as a false signal of microbial generalism.

