## Supplementary Information for "A global survey of host, aquatic, and soil microbiomes reveals shared abundance and genomic features between bacterial and fungal generalists"

Running title: Global patterns and generalist traits in microbiomes

Daniel Loos**[1] Ailton Pereira da Costa Filho**[2,3] Bas E. Dutilh[3,4,5] Amelia E. Barber[2,3,*]
Gianni Panagiotou[1,3,6,*]

[1] Department of Microbiome Dynamics, Leibniz Institute for Natural Product Research and Infection Biology – Hans Knöll Institute. Jena, Germany.
[2] Junior Research Group Fungal Informatics, Institute of Microbiology, Friedrich Schiller University. Jena, Germany.
[3] Cluster of Excellence Balance of the Microverse, Friedrich Schiller University Jena. Jena, Germany.
[4] Institute of Biodiversity, Friedrich Schiller University. Jena, Germany.
[5] Theoretical Biology and Bioinformatics, Utrecht University. Utrecht, the Netherlands.
[6] The University of Hong Kong. Hong Kong SAR, China.

** Equal contribution

### Overview

| Name | Title |
|---|---|
| Supp. Figure 1 | Similar patterns in bacterial and fungal prevalence and abundance across variable rRNA amplicon regions |
| Supp. Figure 2 | Generalists identified are not the result of reagent or sequencing contamination |
| Supp. Figure 3 | Social niche breadth corroborates generalists with significant higher Levins' niche breadth index |
| Supp. Figure 4 | Abundance and co-abundance of generalists and specialists |
| Supp. Figure 5 | Robustness of generalists against varying prevalence |
| Supp. Figure 6 | Node topology metrics in co-abundance networks from host, aquatic, and soil environments |
| Supp. Figure 7 | Specialists have variable impact on alpha diversity |
| Supp. Table 1 | Generalists and specialists identified in this study |

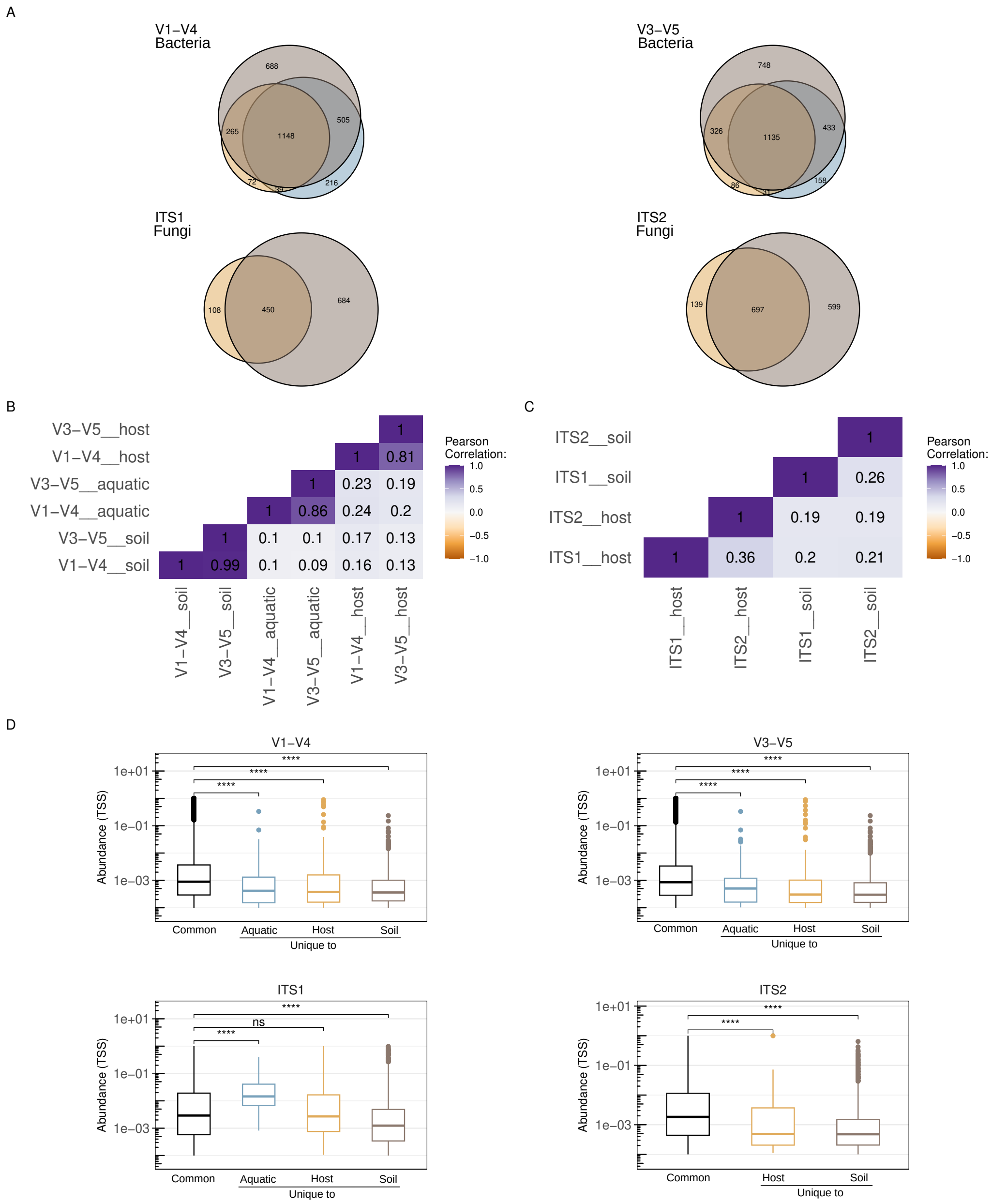


Supplementary Figure 1: **Similar patterns in bacterial and fungal prevalence and abundance across variable rRNA amplicon regions.** (A) Intersection of bacterial and fungal genera found in at least one sample in each environment as Venn diagrams, faceted by region analyzed. The 16S data used in this study was composed of 8 unique primer sets so data was aggregated into studies analyzing sequences within V1-V4 regions and those analyzing sequences the V3-V5 regions. (B-C) Pearson correlation coefficients between rRNA region and environments for bacteria (B) and fungi (C). Significance determined with two-sided t-test, $p < 0.001$ for all correlations. (D) Abundance comparisons of common and unique genera subsetted by rRNA region. A genus was considered present in a sample using a threshold of abundance $> 0.01\%$. Significance determined by Wilcoxon rank sum test; *** denotes $p < 0.001$. No study used ITS2 to capture fungal diversity in aquatic environments, so this comparison is omitted from (C) and (D).

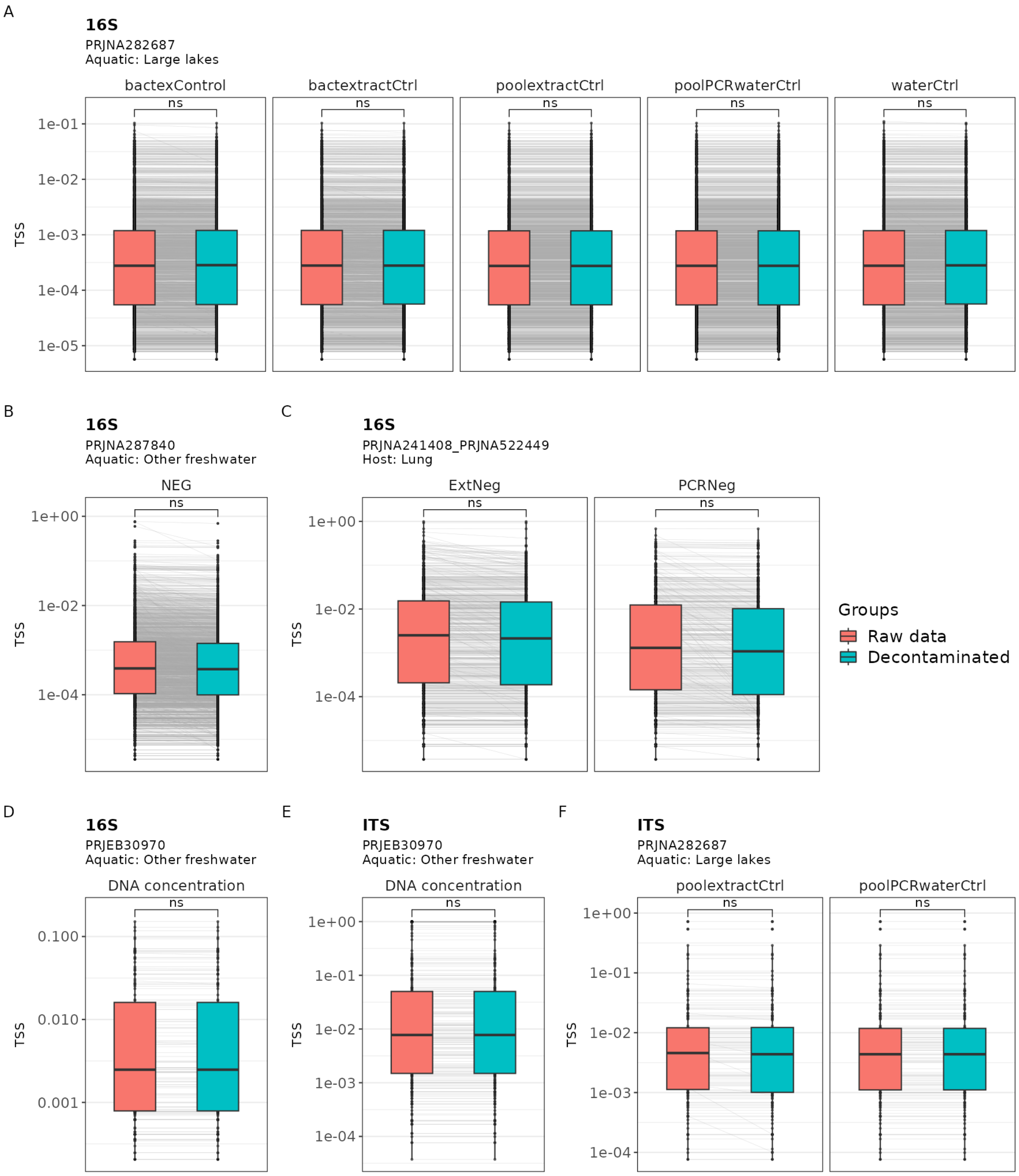


Supplementary Figure 2: **Generalists identified are not the result of reagent or sequencing contamination.** Comparison of the TSS normalized abundance of the generalists identified in the raw data compared to their abundance after decontamination. Connected dots behind the box plots indicate the abundance of a generalist taxa in a sample in the raw data compared to the same taxa in the sample after decontamination. Negative controls sequenced by the project submitter were used in (A-C, F) with decontamination performed using the 'prevalence' mode of the deconam R package. The 'frequency' mode of deconam was used for (D-E), where contaminant OTUs were identified based on their frequency varying inversely with the DNA concentration of the input DNA. Significance determined by Wilcoxon rank sum test; ns denotes $p > 0.05$.

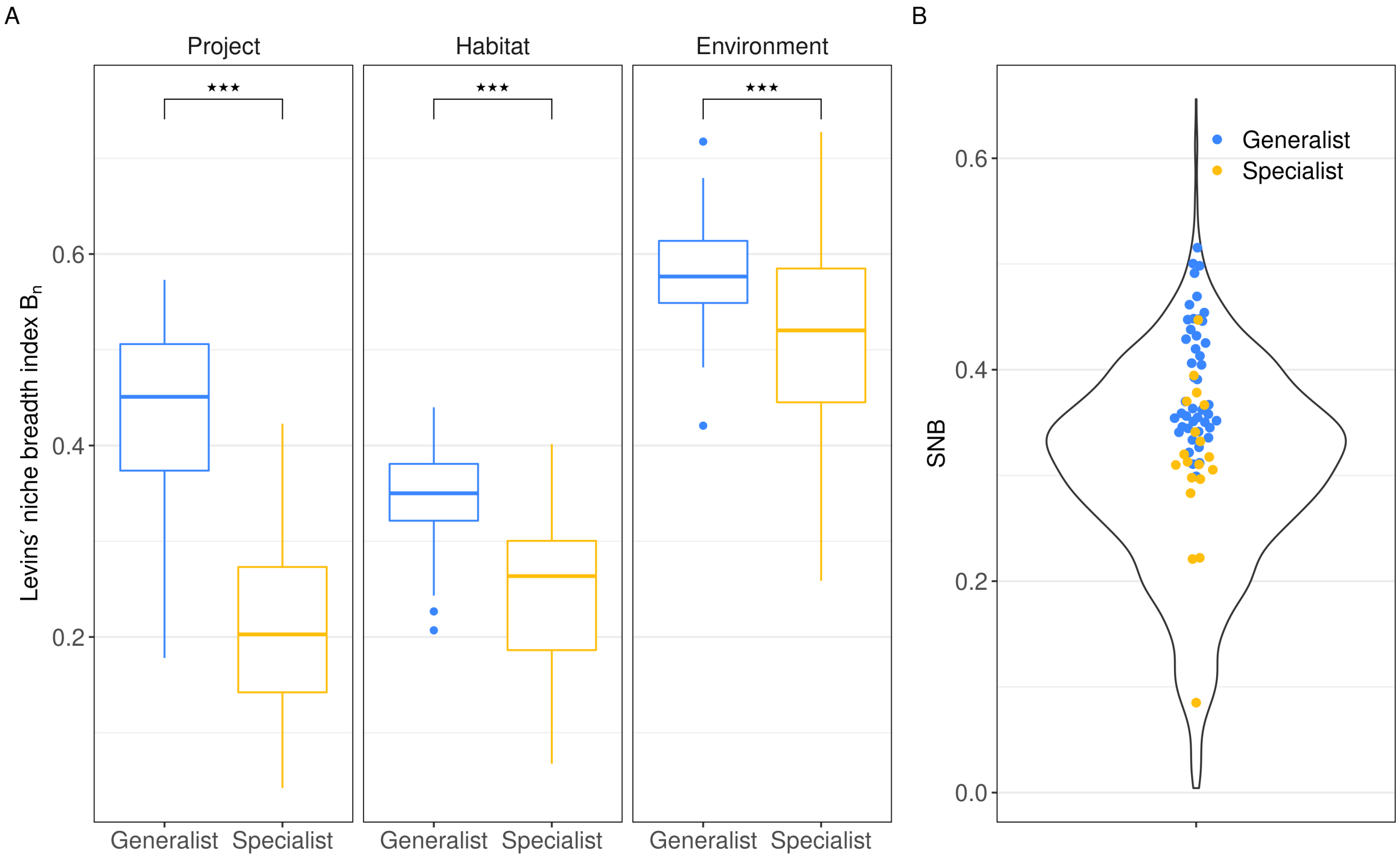


Supplementary Figure 3: **Social niche breadth corroborates generalists with significant higher Levins' niche breadth index.** (A) Levins' niche breadth index (Bn) of generalists and specialists. Generalists are genera found in at least one habitat from each of the three environments (host, soil, aquatic)b with high prevalence (>40%), while specialists are genera with a high prevalence (>40%) in one habitat and low (<5%) prevalence in every other. A relative abundance (RA) cut-off of 0.01% was used for the definition of both generalists and specialists. Significance determined by Wilcoxon rank sum test; *** denotes $p < 0.001$. (B). Social niche breadth score of 3,128 bacterial genera from an independent global microbiome dataset (1) . Generalists and specialists identified in this study are indicated in blue and yellow.

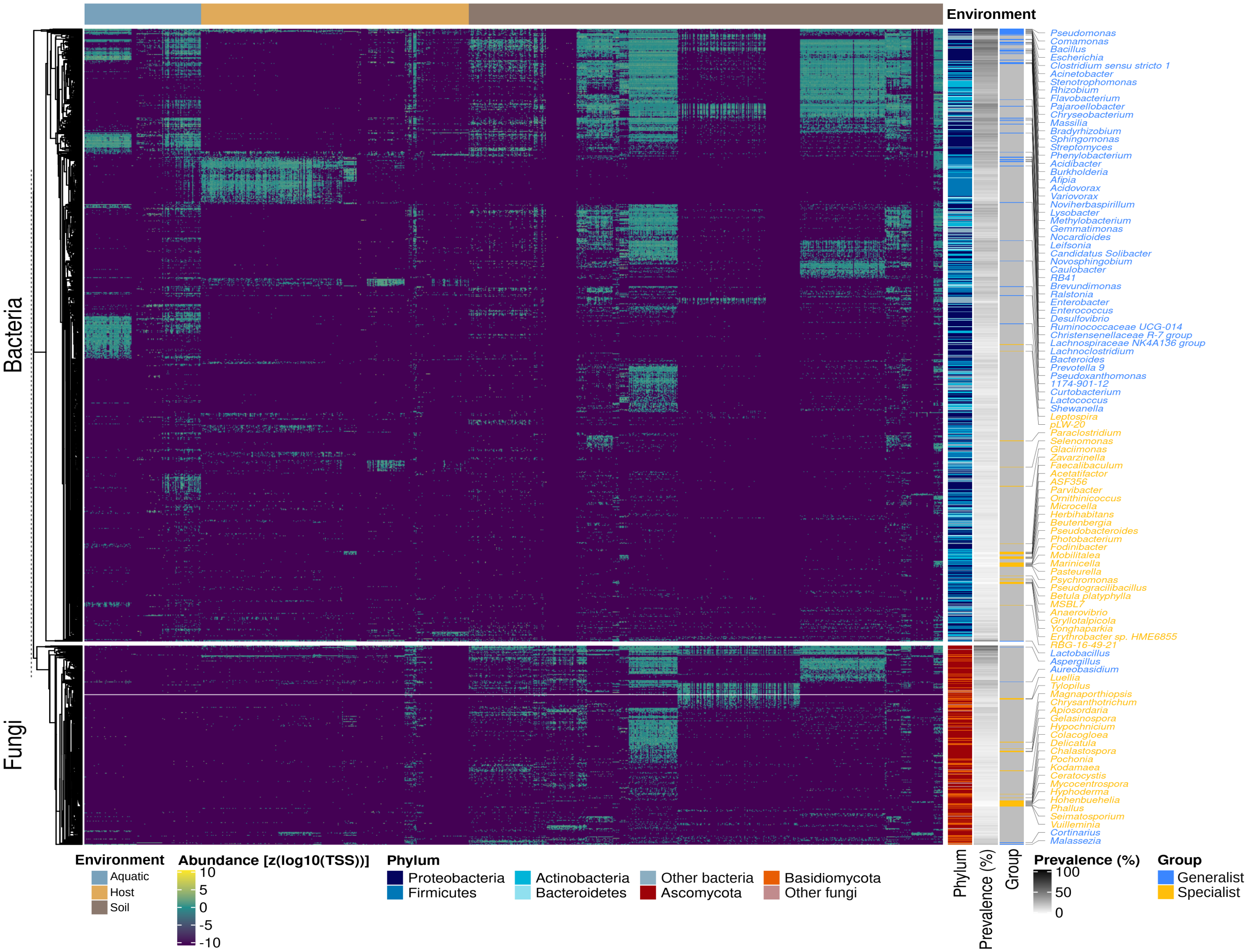


Supplementary Figure 4: **Abundance and co-abundance of generalists and specialists.** Abundance heatmap of bacteria and fungal genera. Samples are arranged by environment (top bar) with only abundance values >0.01% displayed. Right bars indicate phylum and prevalence among all samples, where only taxa with >5% prevalence are displayed. Generalists (>40% prevalence in at least one habitat in all three environments) and specialists (>40% prevalence in one habitat and <5% prevalence in all others) are annotated in blue and yellow, respectively. The dendrogram was calculated using average linkage of Manhattan distances between taxa abundance profiles.

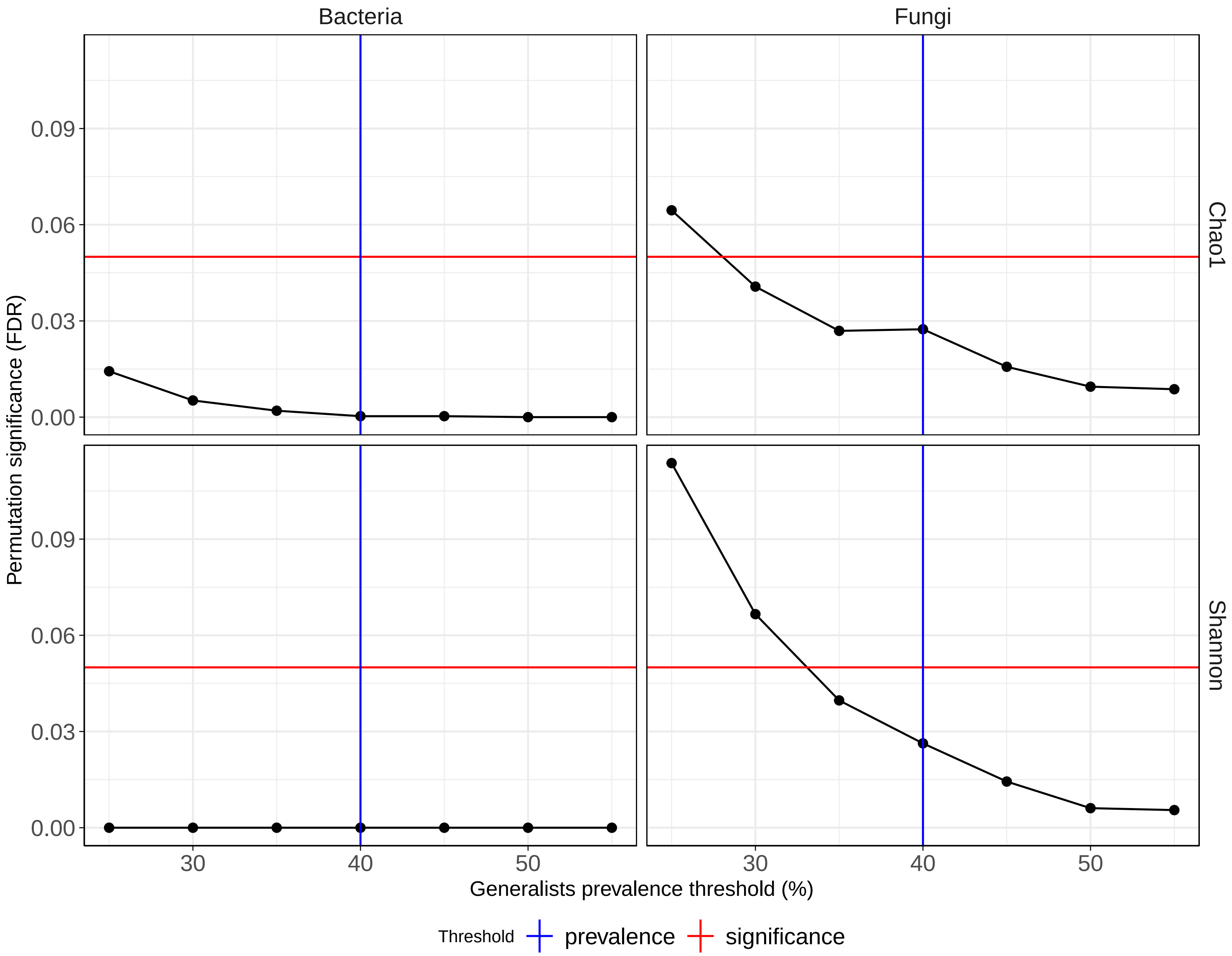


Supplementary Figure 5: **Robustness of generalists against varying prevalence.** Different prevalence thresholds were used to test whether the median alpha diversity in samples without N generalists was lower than in samples without N random taxa. 10,000 times repeated permutation test to sample n taxa without replacement. The test was significant using a wide range of prevalence thresholds around the selected 40%, indicating that this selection is a robust definition of generalists.

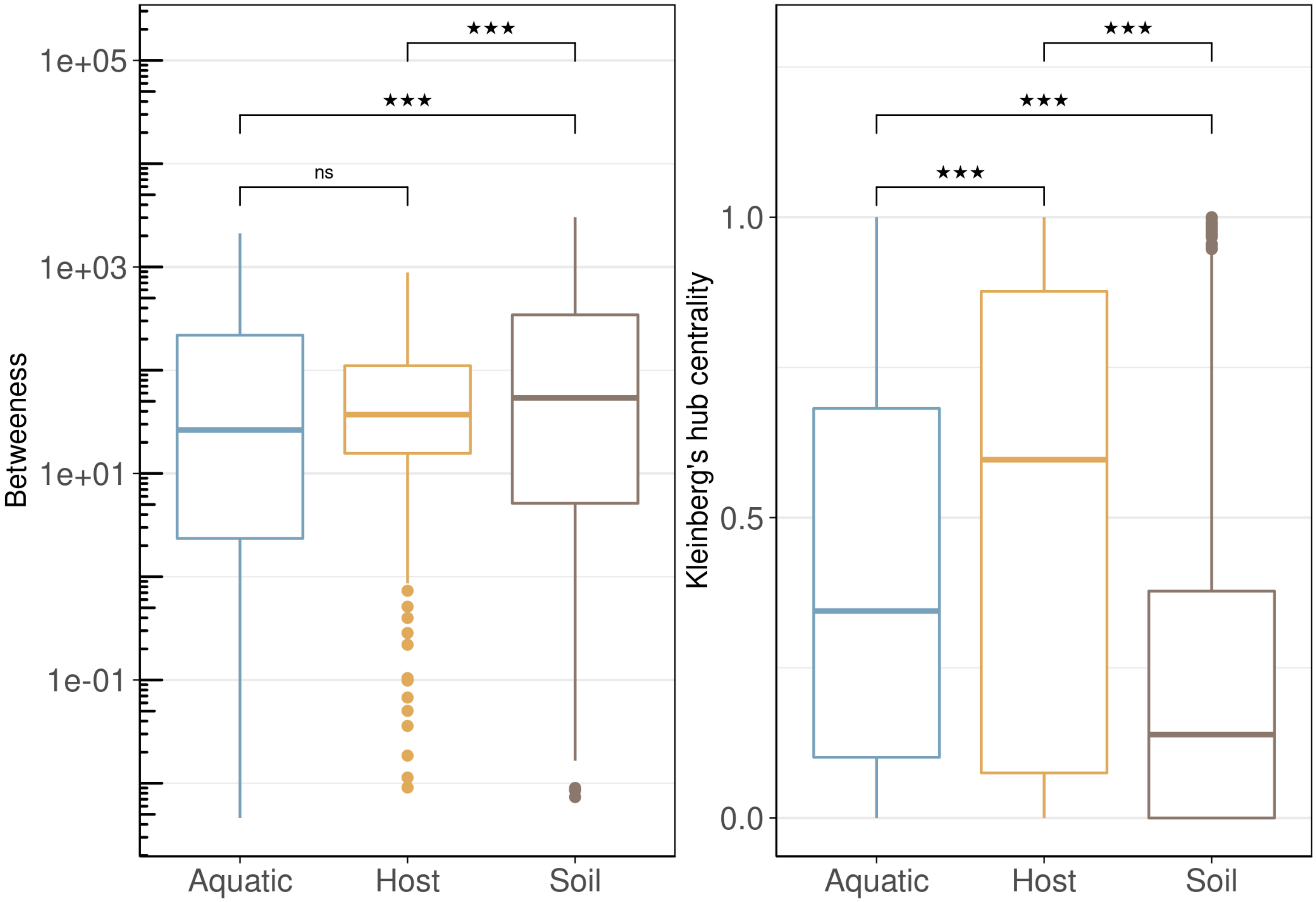


Supplementary Figure 6: **Node topology metrics in co-abundance networks from host, aquatic, and soil environments.** SparCC was used to estimate the correlation between pairs of genera shared between all three environments (N=1,188 bacteria and N=184 fungal genera for input). Networks were generated for each environment individually and edges with a correlation coefficient < 0.2 or an FDR-adjusted p value > 0.05 discarded. Betweenness centrality and Kleinberg's hub centrality scores were calculated for each node (genus) in each network. Significant differences in node scores between each node (genus) in each network. Significant differences in node scores between environments indicates different co-abundance patterns across the three environments (Wilcoxon Rank Sum test, *** denotes p < 0.001).

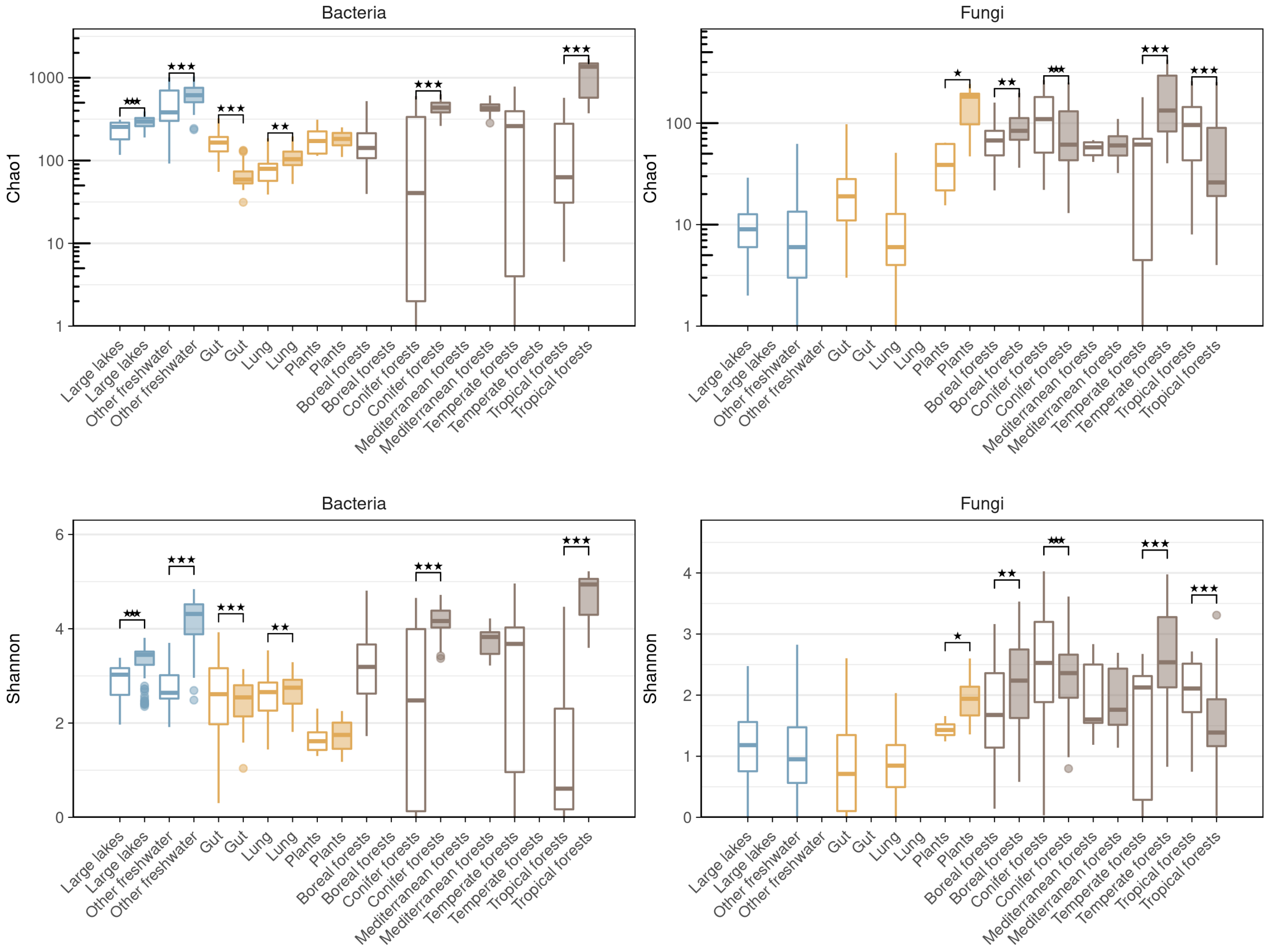


Supplementary Figure 7: **Specialists have variable impact on alpha diversity.** Chao1 and Shannon alpha diversity of bacterial and fungal communities was calculated for all samples. Samples were then pooled by habitat, based on deposited metadata, and further subdivided by whether they contained any specialist (coloured fill) or not (no fill). Some habitats did not contain a specialist so only a single boxplot is visualized. The alpha diversity was higher in some sample group comparisons and lower in others, indicating a complex association between specialists and alpha diversity.

### Supplementary Tables

#### Supplementary Table 1: Generalists and specialists identified in this study.

| Genus | Type | Bioproject | Kingdom |
|---|---|---|---|
| *1174-901-12* | generalist | | Bacteria |
| *Acidibacter* | generalist | | Bacteria |
| *Acidovorax* | generalist | | Bacteria |
| *Acinetobacter* | generalist | | Bacteria |
| *Afipia* | generalist | | Bacteria |
| *Allorhizobium-Neorhizobium-Pararhizobium-Rhizobium* | generalist | | Bacteria |
| *Aspergillus* | generalist | | Fungi |
| *Aureobasidium* | generalist | | Fungi |
| *Bacillus* | generalist | | Bacteria |
| *Bacteroides* | generalist | | Bacteria |
| *Bradyrhizobium* | generalist | | Bacteria |
| *Brevundimonas* | generalist | | Bacteria |
| *Burkholderia-Caballeronia-Paraburkholderia* | generalist | | Bacteria |
| *Candidatus Solibacter* | generalist | | Bacteria |
| *Caulobacter* | generalist | | Bacteria |
| *Christensenellaceae R-7 group* | generalist | | Bacteria |
| *Chryseobacterium* | generalist | | Bacteria |
| *Clostridium sensu stricto 1* | generalist | | Bacteria |
| *Comamonas* | generalist | | Bacteria |
| *Cortinarius* | generalist | | Fungi |
| *Curtobacterium* | generalist | | Bacteria |
| *Desulfovibrio* | generalist | | Bacteria |
| *Enterobacter* | generalist | | Bacteria |

| | | |
|---|---|---|
| *Enterococcus* | generalist | Bacteria |
| *Escherichia-Shigella* | generalist | Bacteria |
| *Flavobacterium* | generalist | Bacteria |
| *Gemmatimonas* | generalist | Bacteria |
| *Lachnoclostridium* | generalist | Bacteria |
| *Lachnospiraceae NK4A136 group* | generalist | Bacteria |
| *Lactobacillus* | generalist | Bacteria |
| *Lactococcus* | generalist | Bacteria |
| *Leifsonia* | generalist | Bacteria |
| *Lysobacter* | generalist | Bacteria |
| *Malassezia* | generalist | Fungi |
| *Massilia* | generalist | Bacteria |
| *Methylobacterium* | generalist | Bacteria |
| *Nocardioides* | generalist | Bacteria |
| *Noviherbaspirillum* | generalist | Bacteria |
| *Novosphingobium* | generalist | Bacteria |
| *Pajaroellobacter* | generalist | Bacteria |
| *Phenylobacterium* | generalist | Bacteria |
| *Prevotella 9* | generalist | Bacteria |
| *Pseudomonas* | generalist | Bacteria |
| *Pseudoxanthomonas* | generalist | Bacteria |
| *Ralstonia* | generalist | Bacteria |
| *RB41* | generalist | Bacteria |
| *Ruminococcaceae UCG-014* | generalist | Bacteria |
| *Shewanella* | generalist | Bacteria |
| *Sphingomonas* | generalist | Bacteria |
| *Stenotrophomonas* | generalist | Bacteria |

| | | | |
|---|---|---|---|
| *Streptomyces* | generalist | | Bacteria |
| *Variovorax* | generalist | | Bacteria |
| *Betula platyphylla* | specialist | PRJEB23282 | Bacteria |
| *Seimatosporium* | specialist | PRJEB23282 | Fungi |
| *Vuilleminia* | specialist | PRJEB23282 | Fungi |
| *Selenomonas* | specialist | PRJNA241408_PRJNA522449 | Bacteria |
| *Hypochnicium* | specialist | PRJNA263505 | Fungi |
| *MSBL7* | specialist | PRJNA271113 | Bacteria |
| *Leptospira* | specialist | PRJNA282687 | Bacteria |
| *pLW-20* | specialist | PRJNA282687 | Bacteria |
| *RBG-16-49-21* | specialist | PRJNA282687 | Bacteria |
| *Blattella germanica (German cockroach)* | specialist | PRJNA287840 | Bacteria |
| *Glaciimonas* | specialist | PRJNA287840 | Bacteria |
| *Apiosordaria* | specialist | PRJNA406830 | Fungi |
| *Chrysanthotrichum* | specialist | PRJNA406830 | Fungi |
| *Colacogloea* | specialist | PRJNA406830 | Fungi |
| *Gelasinospora* | specialist | PRJNA406830 | Fungi |
| *Magnaporthiopsis* | specialist | PRJNA406830 | Fungi |
| *Anaerovibrio* | specialist | PRJNA415280_PRJNA415285 | Bacteria |
| *Erythrobacter sp. HME6855* | specialist | PRJNA415280_PRJNA415285 | Bacteria |
| *Gryllotalpicola* | specialist | PRJNA415280_PRJNA415285 | Bacteria |
| *Hyphoderma* | specialist | PRJNA415280_PRJNA415285 | Fungi |
| *Yonghaparkia* | specialist | PRJNA415280_PRJNA415285 | Bacteria |
| *Mycocentrospora* | specialist | PRJNA418896 | Fungi |
| *Pochonia* | specialist | PRJNA418896 | Fungi |
| *Pseudogracilibacillus* | specialist | PRJNA432446 | Bacteria |
| *Chalastospora* | specialist | PRJNA450848 | Fungi |

| | | | |
|---|---|---|---|
| *Ceratocystis* | specialist | PRJNA492720 | Fungi |
| *Kodamaea* | specialist | PRJNA492720 | Fungi |
| *Acetatifactor* | specialist | PRJNA496065 | Bacteria |
| *ASF356* | specialist | PRJNA496065 | Bacteria |
| *Faecalibaculum* | specialist | PRJNA496065 | Bacteria |
| *Parvibacter* | specialist | PRJNA496065 | Bacteria |
| *Luellia* | specialist | PRJNA517449 | Fungi |
| *Tylopilus* | specialist | PRJNA517449 | Fungi |
| *Fodinibacter* | specialist | PRJNA528359 | Bacteria |
| *Marinicella* | specialist | PRJNA528359 | Bacteria |
| *Mobilitalea* | specialist | PRJNA528359 | Bacteria |
| *Pasteurella* | specialist | PRJNA528359 | Bacteria |
| *Phallus* | specialist | PRJNA528359 | Fungi |
| *Photobacterium* | specialist | PRJNA528359 | Bacteria |
| *Pseudobacteroides* | specialist | PRJNA528359 | Bacteria |
| *Psychromonas* | specialist | PRJNA528359 | Bacteria |
| *Zavarzinella* | specialist | PRJNA550037 | Bacteria |
| *Beutenbergia* | specialist | PRJNA561410_PRJNA561568 | Bacteria |
| *Herbihabitans* | specialist | PRJNA561410_PRJNA561568 | Bacteria |
| *Hohenbuehelia* | specialist | PRJNA561410_PRJNA561568 | Fungi |
| *Microcella* | specialist | PRJNA561410_PRJNA561568 | Bacteria |
| *Ornithinicoccus* | specialist | PRJNA561410_PRJNA561568 | Bacteria |
| *Delicatula* | specialist | PRJNA606949 | Fungi |
| *Paraclostridium* | specialist | PRJNA606949 | Bacteria |
